# Towards genetic modification of plant-parasitic nematodes: delivery of macromolecules to adults and expression of exogenous mRNA in second stage juveniles

**DOI:** 10.1101/2020.07.15.193052

**Authors:** Olaf Kranse, Helen Beasley, Sally Adams, Andre Pires da Silva, Chris Bell, Catherine Lilley, Peter Urwin, David Bird, Eric Miska, Geert Smant, Godelieve Gheysen, John Jones, Mark Viney, Pierre Abad, Thomas R. Maier, Thomas J. Baum, Shahid Siddique, Valerie Williamson, Alper Akay, Sebastian Eves-van den Akker

## Abstract

Plant-parasitic nematodes are a continuing threat to food security, causing an estimated 100 billion USD in crop losses each year. The most problematic are the obligate sedentary endoparasites (primarily root knot nematodes and cyst nematodes). Progress in understanding their biology is held back by a lack of tools for functional genetics: forward genetics is largely restricted to studies of natural variation in populations, and reverse genetics is entirely reliant on RNA interference. There is an expectation that the development of functional genetic tools would accelerate the progress of research on plant-parasitic nematodes, and hence the development of novel control solutions. Here, we develop some of the foundational biology required to deliver a functional genetic tool kit in plant-parasitic nematodes. We characterise the gonads of male *Heterodera schachtii* and *Meloidogyne hapla* in the context of spermatogenesis. We test and optimise various methods for the delivery, expression, and/or detection of exogenous nucleic acids in plant-parasitic nematodes. We demonstrate that delivery of macromolecules to cyst and root knot nematode male germlines is difficult, but possible. Similarly, we demonstrate the delivery of oligonucleotides to root knot nematode gametes. Finally, we develop a transient expression system in plant-parasitic nematodes by demonstrating the delivery and expression of exogenous mRNA encoding various reporter genes throughout the body of *H. schachtii* juveniles using lipofectamine-based transfection. We anticipate these developments to be independently useful, will expedite the development of genetic modification tools for plant-parasitic nematodes, and ultimately catalyze research on a group of nematodes that threaten global food security.

## Introduction

Plant-parasitic nematodes are a continuing threat to food security, causing an estimated 100 billion USD in crop losses each year (Nicol et al. 2011). There are several different plant-parasitic lifestyles across the phylum Nematoda, the most problematic of which are the obligate sedentary-endoparasites (primarily root-knot nematodes and cyst nematodes). Consequently they are some of the most intensely studied (J. T. Jones et al. 2013). A current focus of the research community is to advance our understanding of plant-parasitic nematode biology in sufficient detail to develop novel methods for control. Progress in this aim is held back by a lack of functional genetic tools: forward genetics in the sedentary endoparasites is restricted to the root-knot nematode *Meloidogyne hapla*, and relies on natural variants as the source of mappable polymorphisms (Thomas and Williamson 2013); reverse genetics is entirely reliant on RNA interference (Bakhetia et al. 2005), and is limited by the variable penetrance and stability of the effect. Despite these restrictions, meaningful progress recently has been made. There is nevertheless an expectation that the development of functional genetic tools would accelerate progress in understanding the biology of plant-parasitic nematodes, and thereby also the development of novel control solutions.

There are two major constraints to realising functional genetic tools in the sedentary endoparasitic nematodes: i) lack of knowledge, and ii) biology. Firstly, the development of many functional genetic tools has been in model organisms, and thus builds on a considerable foundation of knowledge that is not yet available for plant-parasitic nematodes. For example, and to the best of our knowledge, no characterized genetic modifications have been identified in plant-parasitic nematodes that give rise to a readily scorable phenotype *in nematoda* (apart from the inability to complete the lifecycle). Secondly, the biology of plant-parasitic obligate sedentary endoparasites is generally not conducive to the technical steps required for genetic modification. Specifically, second stage juveniles (J2) hatch from eggs in the soil. At this stage, the germline in cyst nematodes consists of two non-differentiated germ cell primordia enclosed in two epithelial cap cells, and located approximately 65% of the body length from the anterior end (Subbotin, Mundo-Ocampo, and Baldwin 2010). J2s infect the roots of plants, and cause plant tissue to re-differentiate into a nematode-induced feeding site from which nematodes withdraw all their nutrition. After induction of the feeding site, nematodes become sedentary. Sexual identity is environmentally determined: J2s that induce fully functional feeding sites at an appropriate place to connect to the vascular tissues in the roots become female, while J2 that induce smaller feeding sites, in less favourable locations, become male (Trudgill 1967). In females the germ-cell primordia develops into a didelphic gonad. However, females become opaque, remain attached to the root for their entire life, and their germline is therefore inaccessible. In those juveniles that develop into males, a single gonad branch develops, and the animal regains motility and leaves the root in order to locate and inseminate the sedentary female nematodes. In the case of the sexual (obligate or facultative) sedentary endoparasites, males are therefore the only life stage with a mature germ line that is accessible to manipulation. Their use in hundreds of controlled crosses (Guo et al. 2017) confirms they are fully competent to mate. For obligate parthenogenetic sedentary endoparasites (e.g. the root-knot nematode *Meloidogyne incognita*), males are produced but they do not contribute to the gene pool and, therefore, there is no life stage with a mature germline that is accessible to manipulation. Notwithstanding these challenges, the life cycle of the sedentary endoparasites is at best several weeks (e.g. *Meloidogyne hapla*) and at worst several months (e.g. some *Globodera* have a dormancy period between generations).

Functional genetic tools, such as CRISPR-Cas9 mediated genome editing have been developed in a number of nematode species. These include the widely studied *C. elegans* (Frøkjær-Jensen 2013; Friedland et al. 2013), *C. remanei* (Yin et al. 2018), and *Pristionchus pacificus* (Witte et al. 2015), but also relatively recently in other more challenging (animal) parasitic species (e.g. *Strongyloides* spp., *Auanema freiburgensis* and *A. rhodensis* (Lok 2019; Adams et al. 2019; Gang et al. 2017; O'Halloran 2019)). In this paper we aim to develop some of the foundational biology required to deliver a functional reverse genetics “toolkit” to plant-parasitic nematology. We characterised the germlines of male *Heterodera schachtii* and *M. hapla*. We tested and optimised various methods for the delivery, expression, and/or detection of exogenous nucleic acids in plant-parasitic nematodes. We demonstrate that delivery of macromolecules to cyst and root-knot nematode male germlines is difficult, but possible. Similarly, we are able to deliver oligonucleotides to root-knot nematode germlines. Finally, we show the delivery and expression of exogenous mRNA encoding various reporter genes throughout the body of *H. schachtii* juveniles using lipofectamine-based transfection. Taken together we anticipate these developments to be useful in their own right, expedite the development of genetic modification protocol/s for sedentary endoparasitic nematodes, and ultimately catalyze research on a group of nematodes that threaten global food security.

## Results

### Characterisation of the gonad of adult motile plant-parasitic nematodes

In order to develop a procedure for the genetic modification of plant-parasitic nematodes we need to deliver macromolecules to the germline. Females of sedentary obligate biotrophs (including root-knot and cyst nematodes) are inaccessible. Males however, regain motility to mate with females and therefore, in principle, are accessible for manipulation. We therefore characterised the germlines of male *Heterodera schachtii* and *Meloidogyne hapla* to guide the delivery of macromolecules.

Generally, the gonad of *H. schachtii* and *M. hapla* males is single ended, occupies most of the volume of two thirds to one third of the nematode body length, and appears cellularised, rather than syncytial, throughout. Notwithstanding the considerable variation within species, *M. hapla* males are approximately twice as long and twice as wide as *H. schachtii* males. Most notably, the morphology of germ cells varies considerably between *H. schachtii* and *M. hapla*. The morphology of *H. schachtii* germ cells is uniform from distal tip to proximal end of the germline (Figure 1A). This is consistent with the completion of meiosis prior to the final moult to adult male (Kempton, Clark, and Shepherd 1973). Viewed under DIC-microscopy, the *H. schachtii*germ cells appear as irregular polygons with compact nuclei reminiscent of spermatids in *C. elegans*, and are tightly packed along the entire gonad. Up to four can be found abreast within the gonad of *H. schachtii*. In contrast, germ cells of *M. hapla* vary considerably from the distal tip to the proximal end of the gonad (Figure 1B) as continued sperm development can occur after adult males regain motility (Shepherd and Clark 1983). At the distal tip of the *M. hapla* gonad individual spherical cells are evident, less tightly packed than in *H. schachtii*, and surrounded by a matrix. In the mid-gonad, the cells are larger than at the distal tip and assume an irregular pentagonal shape when viewed under a DIC microscope. More posteriorly, the cells are larger still (only two can be found abreast within the much wider germline of *M. hapla*) and the contents appear granular. These cells resemble spermatocytes of *C. elegans*. Towards the proximal end of the *M. hapla* gonad, the cells are not unlike those found throughout the *H. schachtii* germline: they appear irregular polygons, the nuclei are compact, and they are packed approximately 3-4 abreast in the gonad (therefore approximately twice the size of those in *H. schachtii*). These cells resemble spermatids of *C. elegans* (L’Hernault 2006).

**Figure 1:**
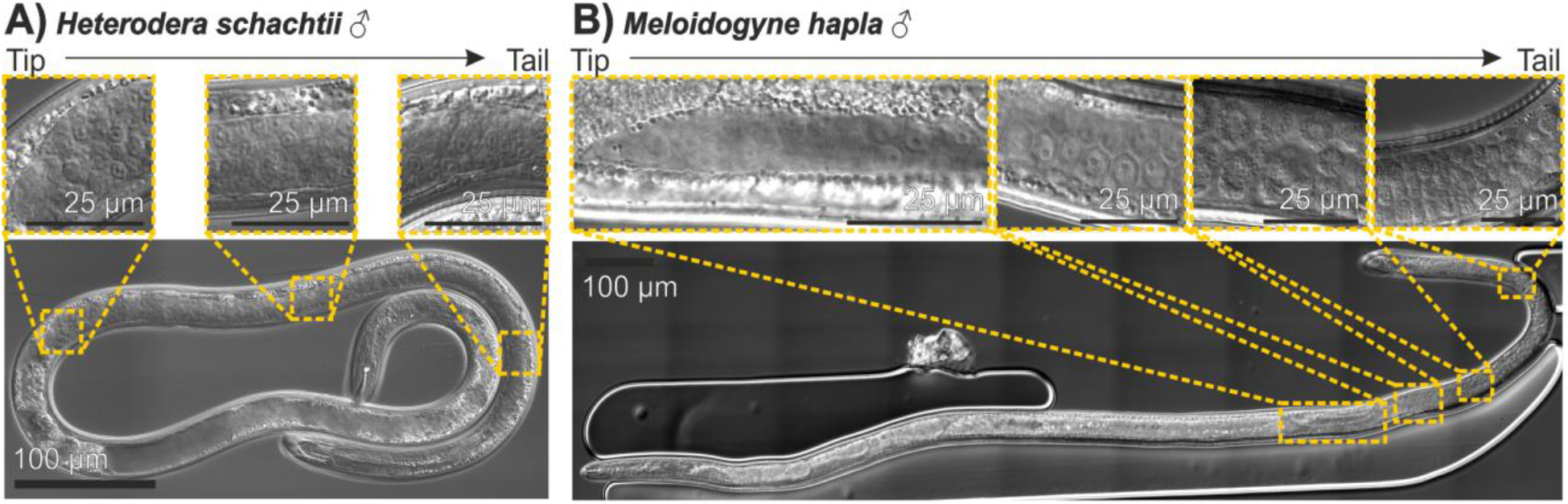
The gonad of cyst and root-knot nematode males. A) *H. schachtii* male. Uniformity of germ cell size and shape from distal tip to proximal end is shown in three inset panels. B) *M. hapla* male. Variation in germ cell size and shape from distal tip to proximal end is shown in four inset panels.

From initial observations we noticed considerable variation in the proportion of the *H. schachtii* male body occupied by the germline. To characterise this phenomenon, we developed a method to collect virgin males at specific times post emergence (Figure 2A). In brief, segments of roots with differentiated males that had not yet emerged, or those dislodged from the root, were collected and stored in PBS at room temperature in a 96 well plate. Adult males that had emerged were removed each day from the 96 well plate, and stored in the absence of females until measured. Males were imaged under DIC microscopy, and measured using FIJI software (e.g. Figure 2A). Notwithstanding large overall variation in absolute body size, the proportion of the body that is occupied by the germline decreases over time from approximately 65%, to approximately 45% (Figure 2B, n = 7-9, Mann-Whitney U, p<0.05). The decrease in the proportionate length of the gonad is likely a combination of more modest but individually not significant increases in body length and decreases in gonad length (Figure 2C and D).

**Figure 2:**
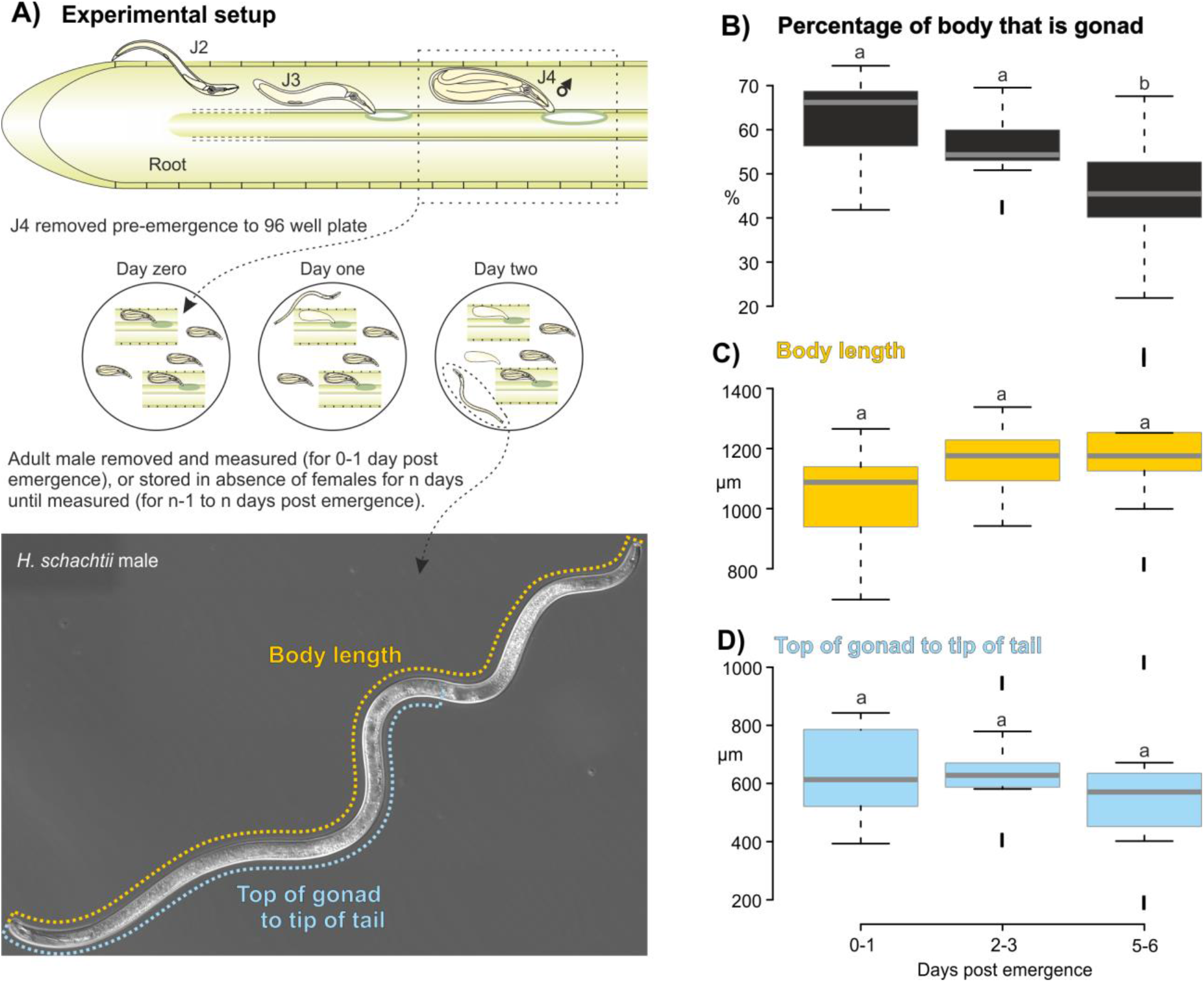
Measuring the change in *H. schachtii* male gonad over time. A) Schematic representation of experimental setup to collect differentiated males pre-emergence with representative measurements of *H. schachtii* males and germline. B-D) comparison of germline and body length at 0-1, 2-3, and 5-6 days post emergence of virgin males showing percentage of body length occupied by germline, body length, and gonad length respectively. Horizontal line in box plot represents median value, whiskers extend to data points that are less than 1.5 x IQR away from 1st/3rd quartile. Different lowercase letters indicate significant differences at p < 0.05 (Mann-Whitney U test).

As the gonad becomes proportionally shorter, we observed the progressive appearance of large vacuous structures between the head and the distal tip of the gonad in virgin males, with this starting almost immediately after emergence (Figure S1). Male nematodes (presumably some non-virgin) with large vacuous structures can be recovered from infected roots, and so we reason this is not a function of the buffer they were stored in prior to imaging, but rather a natural phenomenon.

**Figure S1:**
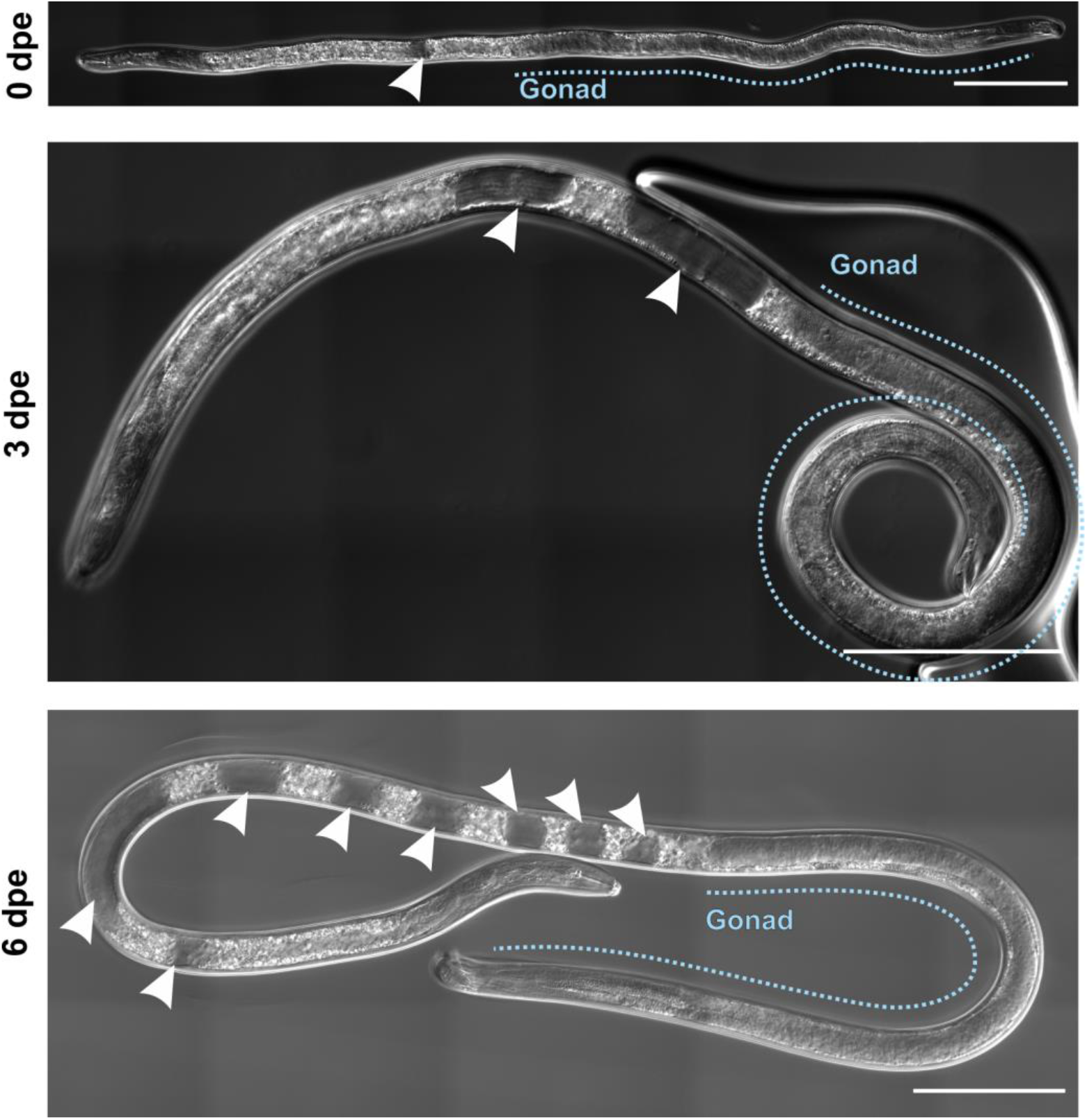
Progressive appearance of large vacuous structures over time in *H. schachtii* virgin males. Representative images of virgin males at 0, 3, and 6 days post emergence (dpe). Arrows indicate vacuous structures. Blue line indicates the region of the body occupied by the gonad. Scale bars indicate 100 μm.

### Delivery of macromolecules to male gonads

Initially, we followed the microinjection methods used for the transformation of *C. elegans* (Evans 2006). We prepared dried 2 % w/v agarose pads and used the Halocarbon Oil 700 for the immobilisation of the nematodes. *H. schachtii* was not immobilised using this method. To overcome this problem we used dried 5 % w/v agarose pads that improved the immobilisation, but still to a much lesser extent than is achieved with *C. elegans* on dried 2 % w/v agarose pads. Our initial attempts using the pre-pulled Eppendorf needles failed mostly due to needle tips breaking when pressed against the hard cuticle of *H. schachtii* or because the needle became blocked by the contents of the nematode flowing into the needle. Using self-pulled needles we were able to insert the intact needle through the cuticle more reliably. However, often this led to bursting of the animal through the injection site. This suggests that *H. schachtii* likely has much higher internal pressure than does *C. elegans*, which leads to bursting upon entry of a needle. In addition, and perhaps related to either high internal pressure and/or stiff body wall, we had to use the highest pressure setting on the Eppendorf Femtojet pump to be able to deliver any material. Even at these maximum pressure settings we often failed to see any delivery from the needle (as judged by visible expansion inside the animal). On a few occasions, inserting the needle into the large vacuous structures (Figure S1) caused them to “burst”, leak their contents, and led to the expansion of the germline. A secondary injection in the gonad was subsequently easier. Nevertheless, upon withdrawal of the needle animals usually leaked through the puncture hole/s. The vacuous structures might contribute to the high internal pressures.

*Meloidogyne hapla* worms, on the other hand, were easier to immobilise using both dried 2 and 5 % w/v agarose pads. Their large size (approximately twice that of *H. schachtii* males) makes identification of the gonad easier under the injection microscope. Using self-pulled needles we were successful in inserting the needle into the gonad and observing the delivery of material through the needle. *Meloidogyne hapla* animals withstood the injection better than *H. schachtii*, and we observed much less bursting despite using the maximum pressure settings. Many animals were mobile in the buffer 2-3 days post injection.

Despite the difficulties, following optimisation, we were able to deliver a membrane permeable DNA dye (Hoechst) to both *H. schachtii* and *M. hapla* male gonad (Fig. 3). In both cases, the dye remains local to the site of injection, is taken up by cells, and stains the nuclei. The observed number and size of fluorescent foci is characteristic of each species’ germline (*H. schachtii* small and numerous, *M. hapla* fewer, larger cells with less dense nuclei).

**Figure 3:**
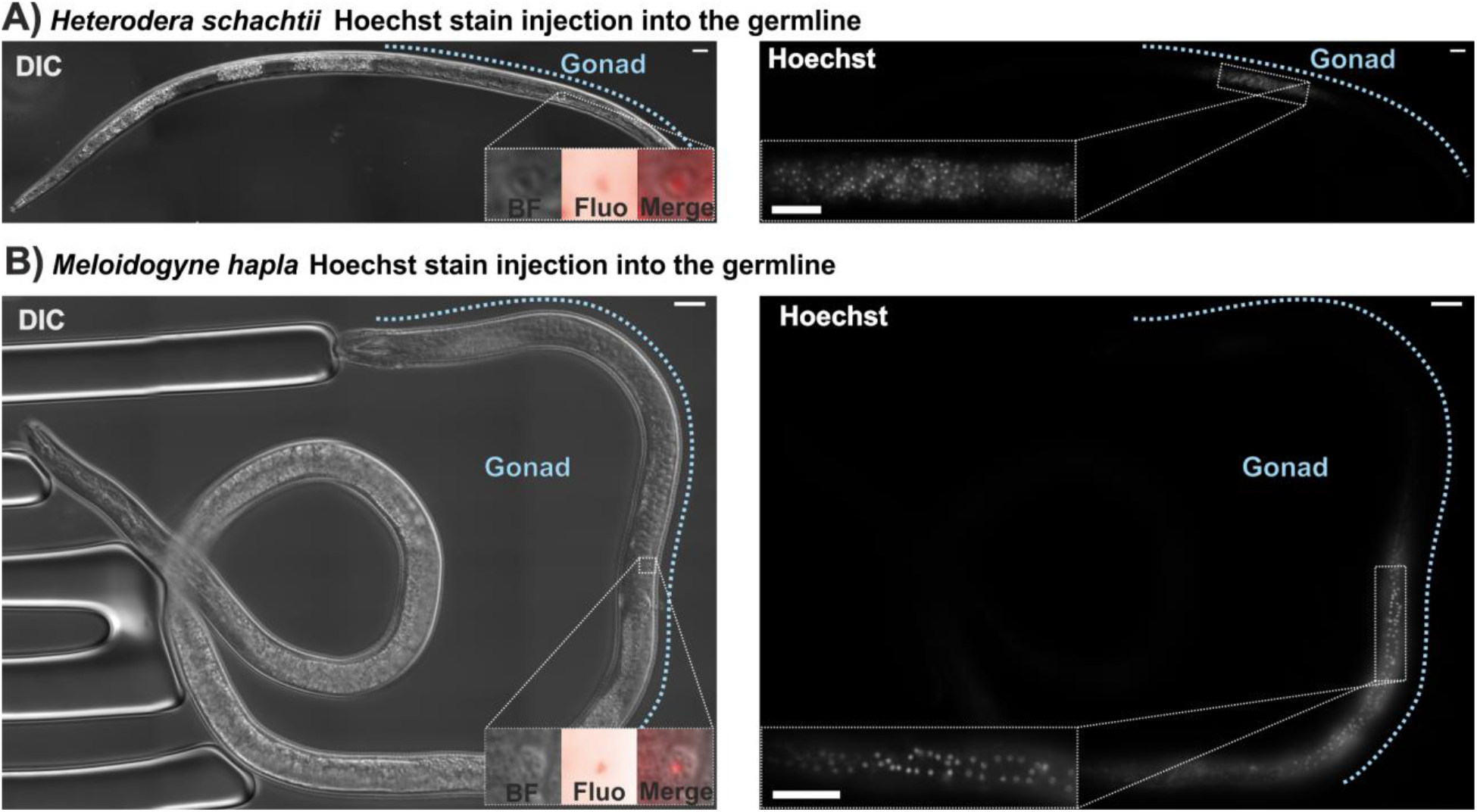
Microinjection of fluorescent dye to adult male gonad of *H. schachtii* and *M. hapla*. Brightfield left, fluorescence right. Gonads indicated by blue dotted lines. Inset in the left panel shows a digital zoom of a single cell within the gonad (brightfield (BF), fluorescence (Fluo), and overlay (Merge)). Inset in the right panel shows a digital zoom of a region of the gonad. A) *H. schachtii* male. B) *M. hapla* male. Scale bars indicate 20 μm.

To further explore macromolecule delivery to the more accessible *M. hapla* gonad, oligonucleotides (Table S1, 63 nt) were synthesised with the covalent addition of 5’ Cy5.5 (excitation 675 nm, emission 694 nm) or FITC (excitation 495 nm, emission 519 nm). Examples of apparently successful and unsuccessful injection of Cy5.5-tagged oligo to the *M. hapla* germline are shown in Figure 4A and B respectively (determined by visible expansion of the worm following injection and detection of abundant fluorescence in the injection site). Similar to injection of Hoechst, the material remains local to the site of injection. In a successful injection the majority of the oligo remains in the space between cells in the germline, and highlights their characteristic shape (*cf* Figure 1B). For most successful injections one, or very few, individual cells proximal to the injection site were extremely bright, perhaps indicative of uptake (Figure 4A).

**Figure 4:**
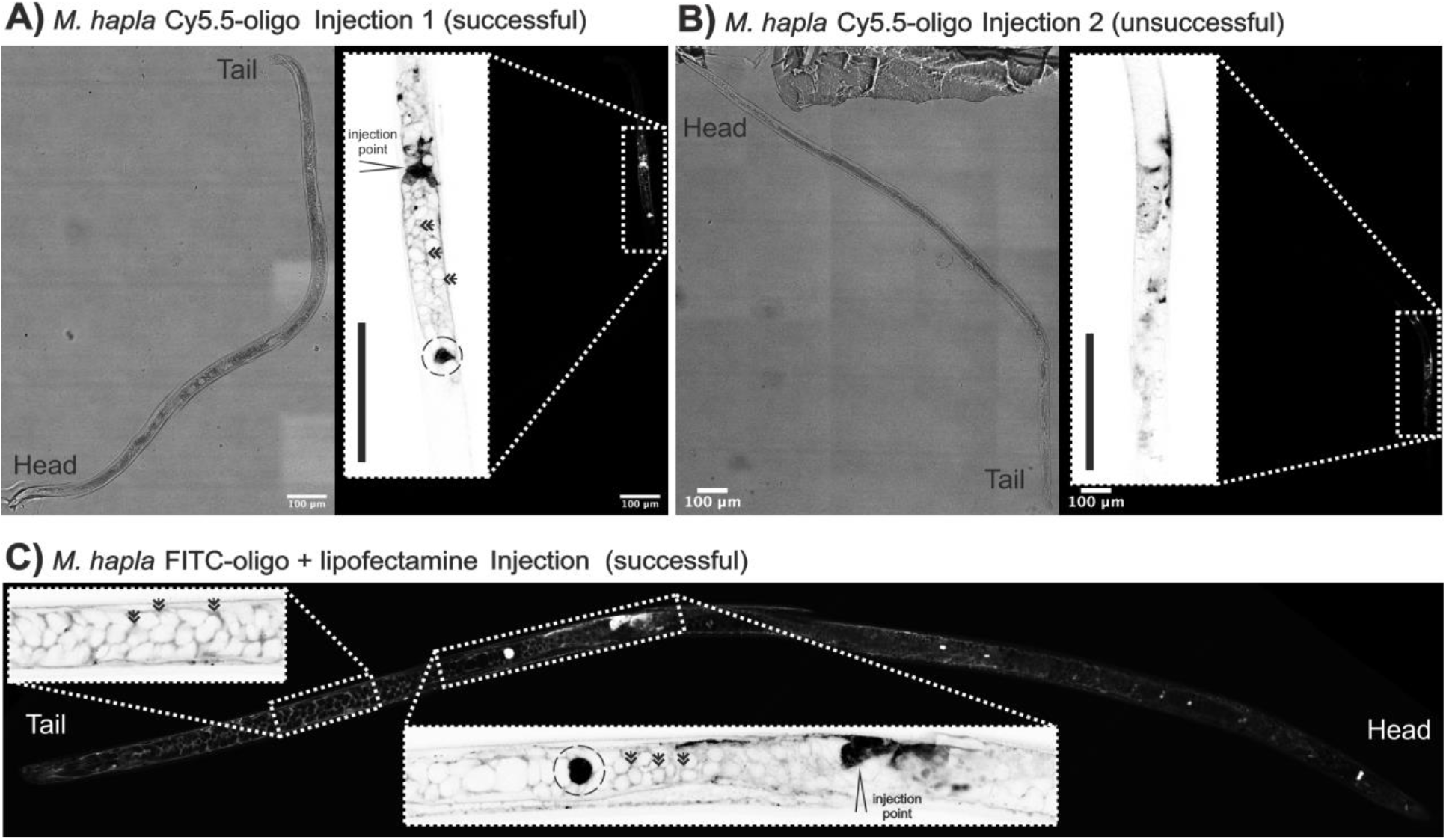
Microinjection of fluorescent-tagged oligonucleotides to the *M. hapla* male germline. A) and B) successful and unsuccessful injection of oligonucleotides-tagged with Cy5.5. Brightfield left, fluorescence right (inset inverted high magnification image of injection site (marked for successful injection). C) Fluorescence image of successful injection of oligonucleotides-tagged with FITC (inset inverted high magnification image of germline tail end (left) and successful injection site (right, marked). Double arrows indicate most fluorescent-tagged-oligonucleotides collect in the space around cells. An example of extremely bright cells, perhaps indicative of uptake, are highlighted with broken circles. Scale bars indicate 100 μm.

We explored the use of liposomes to facilitate delivery of macromolecules. Liposomes are vesicular lipid bilayers that have long been used to deliver various cargo to cells by transfection (Felgner et al. 1987), and more recently to aid microinjection (Adams et al. 2019). We therefore encapsulated FITC-oligo into liposomes (CRISPRMAX lipofectamine) before injection, and injected into the gonad of male *M. hapla* (Figure 4C). Liposomes are vesicular lipid bilayers that have long been used to deliver various cargo to cells by transfection (Felgner et al. 1987), and more recently to aid microinjection (Adams et al. 2019). The resulting fluorescence does not remain local to the site of injection, unlike the injection of the Cy5.5 oligo. However, similar to injection of the Cy5.5 oligo, the majority of the fluorescence remains in the space between cells in the germline, and highlights their characteristic shape. Occasionally, individual cells within the germline, proximal to the injection site, are extremely bright, perhaps similarly indicative of uptake (Figure 4C). This is different in appearance to the injection of Hoechst (Figure 3) because Hoechst is a cell permeable dye that specifically stains nuclei (minimal fluorescence in the cytoplasm but becomes highly fluorescent when bound to DNA) while the oligos are fluorescent in solution and therefore also in the cytoplasm.

### Lipofection-based delivery and expression of mRNA in *H. schachtii* second stage juveniles

Given that *H. schachtii* males are extremely challenging to inject, we developed an alternative lipofection-based method to deliver macromolecules to J2s. mRNAs encoding enhanced GFP (eGFP) were packaged into liposomes and delivered to *H. schachtii* J2 by *in vitro* soaking for 24 hours. Soaked nematodes were washed and imaged using confocal microscopy. On an inverted grayscale, nematodes soaked with mRNA encoding eGFP encapsulated in liposomes were qualitatively darker (i.e. have more fluorescence) than those soaked in lipofectamine alone. However, fluorescence exists in all imaged nematodes (autofluorescence), particularly in the intestine. A quantitative approach was developed to differentiate between eGFP fluorescence and autofluorescence. In brief: greyscale fluorescence images of individual nematodes were extracted, inverted, and the contrast of all cropped images adjusted. Pixels exceeding an empirically derived brightness threshold (see methods for details) were counted and marked (see methods and Figure S2).

**Figure S2:**
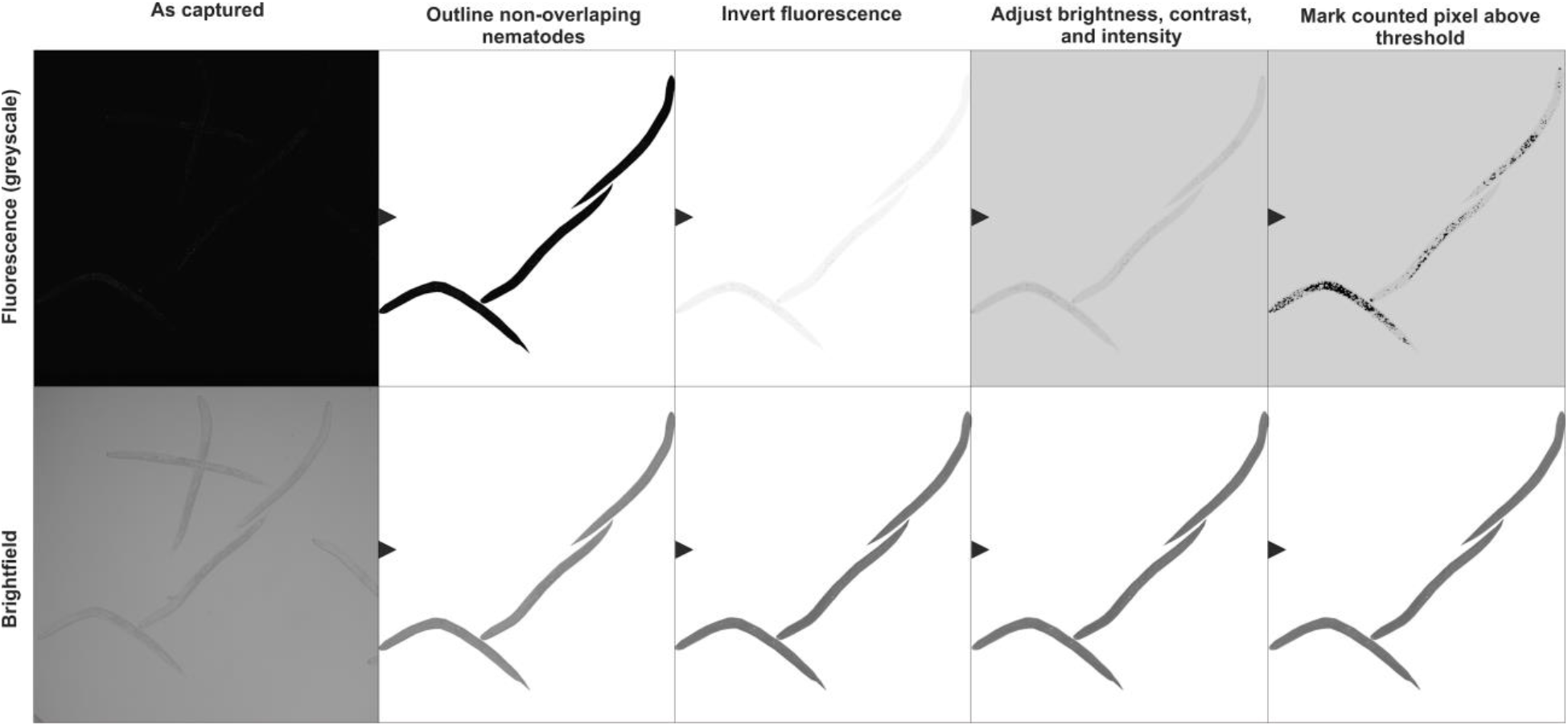
Quantification of fluorescence. To quantify the fluorescence of individual nematodes, the image of each nematode was cropped out, the colours inverted, their brightness, contrast, and intensity adjusted for both treatment and control (Brightness −17%, Contrast + 71%, Intensity −27%), and the number of pixels per nematode that exceeded an empirically determine number of shades of variance (ranging between 0 - 255) from the background grey were counted and marked on the image using a custom script.

Using this method, the intensity of fluorescence between mRNA soaked worms and control soaked worms was quantified and compared. From the representative images in Figure 5A, a clear difference was observed in the fluorescence intensity between the control and mRNA soaked worms. Nematodes soaked with lipofectamine RNAIMAX containing mRNA encoding eGFP were on average six times brighter than lipofectamine-only soaked worms (n = 17, Mann– Whitney U p= 8.57*10^-10^). Additionally, the brightest nematode in the negative control was less bright than the dimmest nematode in the eGFP mRNA treatment using the same microscope settings. From the marked pixels, we can see that: 1) Most autofluorescence in control nematodes is, as expected, restricted to the digestive system of the nematode; and 2) the observed increase in fluorescence in treated nematodes is not just in the digestive system of the nematode, but also throughout the body. In an independent experiment, a comparison was made between the efficacy of two different types of lipofectamine, RNAIMAX and CRISPRMAX (Figure 5A). The RNAIMAX outperformed CRISPRMAX in both brightness and spread of the fluorescence (n = 7 and 9 respectively, Mann–Whitney U p = 3.50*10^-4^).

**Figure 5:**
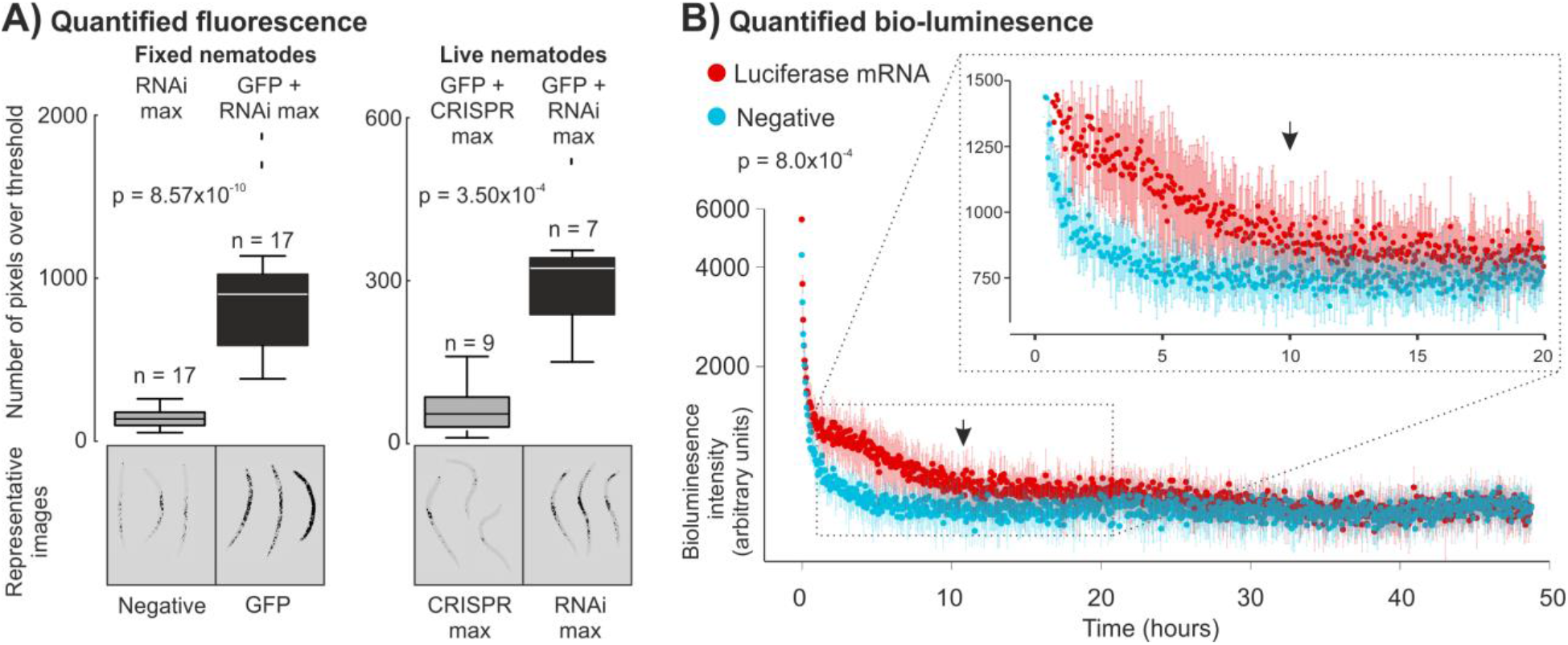
Expression of exogenous mRNAs in second stage juvenile *H. schachtii*. A) Quantification of fluorescence. Representative images of negative control and treated nematodes. Pixels above threshold marked (black) and counts are shown in the boxplot. Horizontal line in box plot represents median value, whiskers extend to data points that are less than 1.5 x IQR away from 1st/3rd quartile. Left, nematodes fixed post lipofection comparing nematodes soaked in empty liposomes (negative) with nematodes soaked in liposomes containing mRNA encoding GFP. Right, live nematodes comparing mRNA encoding GFP encapsulated in either CRISPRMAX or RNAiMAX lipofectamine. P-values are indicated for independent 2-group Mann–Whitney U test. B) Quantified bio-luminescence (arbitrary units plotted on log scale) of live nematodes soaked in mRNA encoding luciferase encapsulated in liposomes (red), or nematodes soaked in empty liposomes (turquoise), measured every 176 seconds for 48.84 hours. Inset, a zoom in of hours 0-20. Arrow indicated 10 hours. The half-life in treated nematodes is compared to the control using the independent 2-group Mann–Whitney U test. Error bars indicate standard deviation of the mean (n = 8) at each time point.

To determine the potential lifetime of transient expression, we repeated the experiment with mRNA encoding luciferase encapsulated in RNAIMAX liposomes and measured nematode light emission every 176 seconds for two days using a CLARIOstar plate reader. Nematodes continued to emit light above background noise luminescence for at least 10 hours, and possibly up to 30 hours (Figure 5B), beyond which the treated and control nematodes were indistinguishable, presumably as substrate, mRNA, or both, are consumed. The decrease in luminescence as a function of time of treated and control nematodes was compared using a Mann-Whitney U test for unpaired data (p =8.0*10^-4^).

### Challenges associated with lipofection-based delivery of CRISPR-Cas components to second stage juvenile *H. schachtii*

Finally, we explored the possibility of using CRISPR-Cas9 to initiate homology directed repair (HDR) and/or non-homologous end joining (NHEJ) in somatic cells of *H. schachtii* juveniles using lipofection. Initially we encapsulated CRISPR-Cas9 protein, guide RNAs, and a single stranded donor DNA fragment designed to introduce an amino acid mutation into the coding sequence of a FAR-1-like gene of *H. schachtii* into CRISPRMAX liposomes. These components were delivered to *H. schachtii* J2s by *in vitro* soaking (Figure 6A). Two negative controls were used: one omitting the guide RNAs but keeping the donor fragment, and one omitting all CRISPR components. We extracted DNA from approximately 20,000 treated nematodes, digested single stranded DNA, and amplified a 429 bp region of interest by PCR using oligonucleotide primers F1:R1, and sequenced the purified amplicons using Illumina technology. In nematodes transfected with donor DNA fragments the expected edit was present in the amplicon, even in the absence of the guide RNAs (Figure 6B). The most parsimonious explanation for this is template switching of the polymerase during amplification from the genome, to the donor DNA fragment, and back. We confirmed this activity *in vitro* by using a second round of PCR, on the purified product of F1:R1, with an oligonucleotide primer F2, that is specific to the desired edit (Figure 6C). Given the promiscuity of template switching, future experiments did not use donor fragments, and focused on detection of NHEJ.

**Figure 6:**
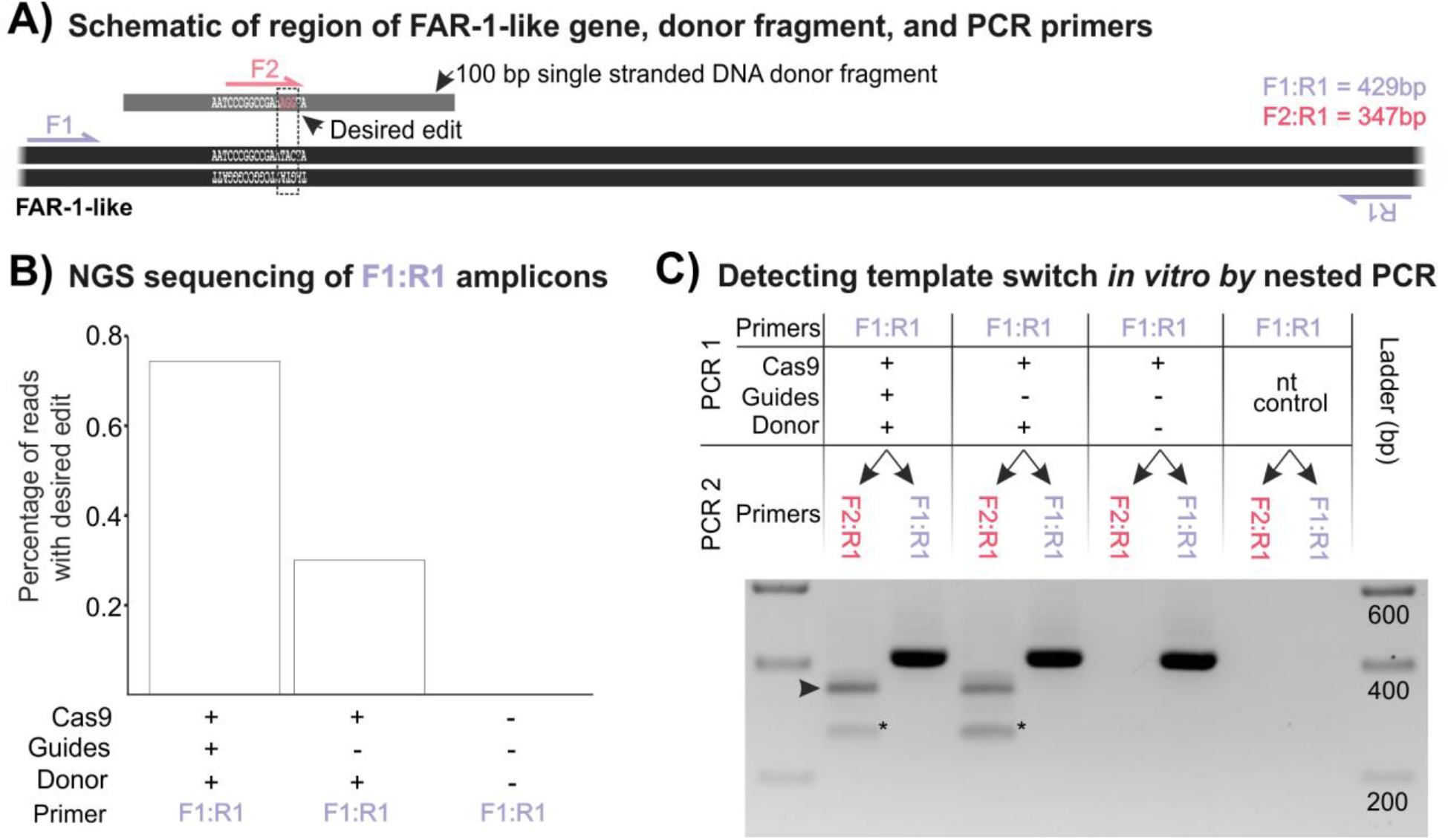
PCR-derived template switching shrouds detection of HDR. A) Schematic representation of a region of FAR-1-like gene of *H. schachtii* indicating primer binding sites and single stranded donor template carrying desired edit (pink bases, AGG from TAC). B) Next Generation Sequencing (NGS) of the F1:R1 amplicon from transfected J2s reveals apparent HDR events, even in the absence of guide RNAs. C) Template switching can be detected in vitro (arrow) using a second primer (F2), on the purified product of F1:R1, that is specific to the desired edit. Star indicates a non-specific amplicon.

Subsequently, twenty eight guide RNAs were designed to target ten genes (at least one, and up to four, guide RNAs per gene (Table S1)). For half of the genes we used guide RNAs modified for increased editing efficiency (2’-O-Methyl at first 3 and last bases, 3’phosphorothiate bonds between first 3 and last 2 bases, Synthego) and five genes used standard guide RNAs (Synthego). Primers were designed to amplify a 300-500 bp fragment that contained the regions targeted by the guide RNAs for each gene (Table S1). For each gene (DRYAD accession doi:10.5061/dryad.r4xgxd296), ribonucleoprotein complexes were assembled and encapsulated in both RNAIMAX and CRISPRMAX liposomes, and independently delivered to juveniles as described for mRNA. DNA was extracted from transfected nematodes, the region of interest amplified, and the amplicon sequenced using 250 bp paired end Illumina reads. All reads were compared to a reference sequence to identify the presence of indels (Figure 7) and substitutions (Figure S3) within the region targeted by each guide RNA (i.e. 1 to 6 bp upstream of the protospacer adjacent motif). Across the data set, there are generally low numbers of reads containing indels and/or substitutions within the guide region/s, and are only two consistent differences between treated and negative control nematodes (albeit with very low number of absolute read support, Gene 1 guide 1 and Gene 8 guide 1). This suggests that either the CRISPR-Cas9 is inducing NHEJ below the detection limit of this experiment and is shrouded by noise (of PCR and/or sequencing) or the CRISPR-Cas9 reagents are not delivered in sufficient quantity/at all to induce NHEJ.

**Figure 7:**
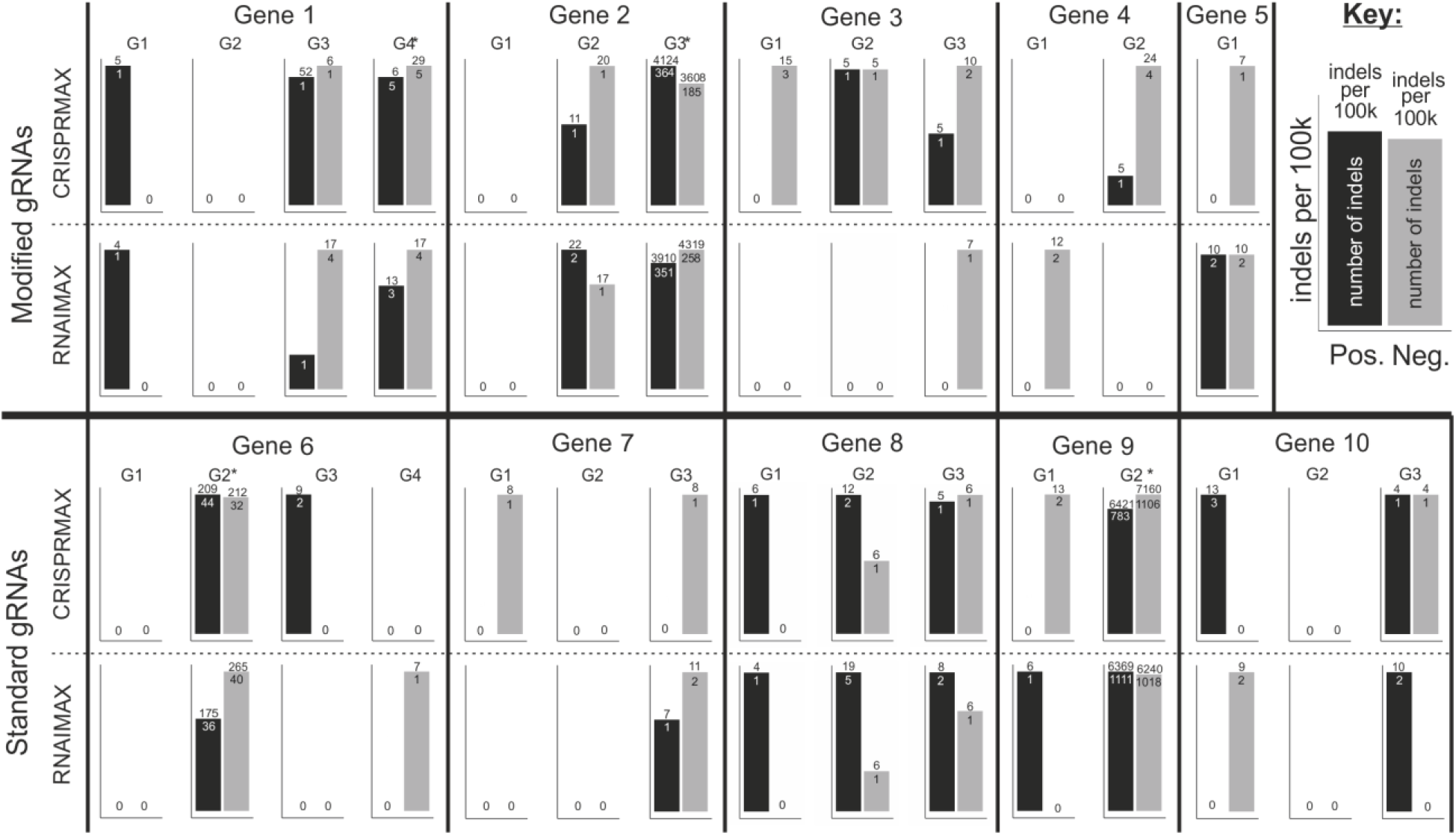
Next generation sequence indel analysis of lipofection-based CRISPR-Cas9 trial. Two lipofection reagents, CRISPRMAX and RNAIMAX were used to deliver CRISPR-cas9 components to *H. schachtii*, guided by either modified (top) or standard (bottom) gRNAs (G1-4). For each of 10 genes, bar graphs show the number of indels per 100,000 reads (number above the bar). Absolute number of reads containing indels is shown within each bar for positive (black bars) or negative (grey bars, omission of guides). Guide regions with several polynucleotides, and therefore high sequencing inaccuracy, are indicated with a *.

**Figure S3:**
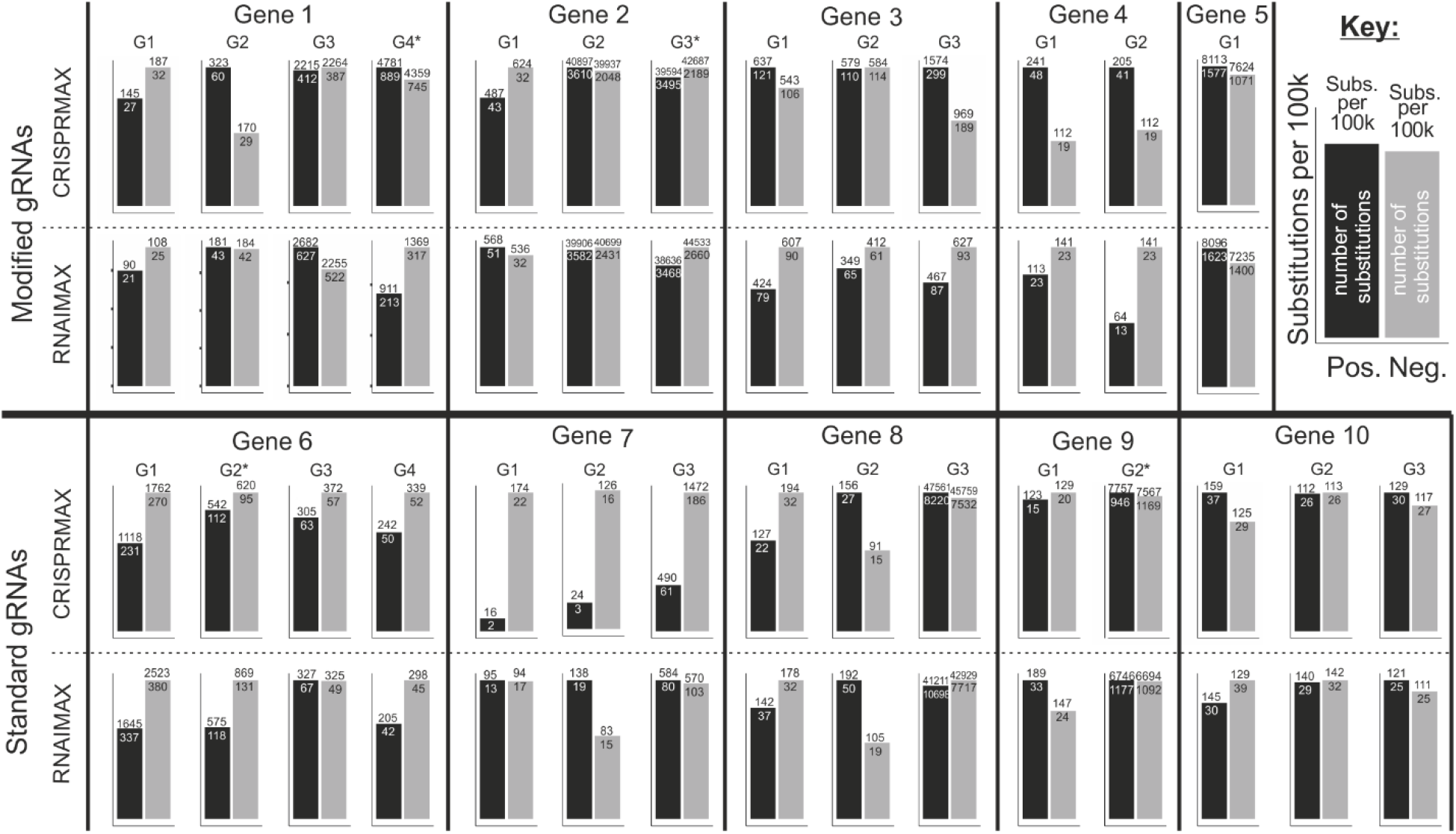
Next generation sequence substitution analysis of lipofection-based CRISPR-Cas9 trial. Two lipofection reagents, CRISPRMAX and RNAIMAX were used to deliver CRISPR-cas9 components to *H. schachtii*, guided by either modified (top) or standard (bottom) gRNAs (G1-4). For each of 10 genes, bar graphs show the number of substitutions per 100,000 reads (number above the bar). Absolute number of reads containing substitutions is shown within each bar for positive (black bars) or negative (grey bars, omission of guides). Guide regions with several polynucleotides, and therefore high sequencing inaccuracy, are indicated with a *.

## Discussion

Obligate sedentary endoparasites are the major contributor to the worldwide crop losses caused by plant-parasitic nematodes, and so are intensely studied. Developing a system to genetically transform them should therefore be top priority, but their biology makes them one of the most difficult species in which to achieve this.

The biology and gonad accessibility are two major barriers to transformation of sedentary endoparasitic plant parasites: the germlines are either not developed enough (juvenile), inaccessible (female), or apparently non-syncytial when accessible (males). Most successful examples of using microinjection to transform nematodes are on species with accessible syncytial gonads (e.g. *Caenorhabditis* spp., *Pristionchus* spp., and the animal parasite *Strongyloides* spp.). One exception may be *Auanema s*pp., although it is not entirely clear whether the gonad is syncytial or not (Adams et al. 2019). In *H. schachtii* and *M. hapla*, males are the only accessible life stage with a developed germ line (albeit non-syncytial). In general male root-knot and cyst nematodes are technically hard to inject: their cuticles are hard, their body is non-elastic, and they appear to be under high internal pressure so that delivering contents from the needle is difficult and unreliable. *Heterodera schachtii* males are less suitable for microinjection than *M. hapla* males, because the former are smaller and their germline cells are fully differentiated, with all cell divisions occurring before the final moult and only spermatids and spermatozoa present in the adult gonad (Kempton, Clark, and Shepherd 1973) and they are less able to accommodate injections. In contrast, all stages of sperm development are present in the adult male gonad of *M. hapla*, from spermatogonia at the tip to mature spermatozoa at the proximal end (Shepherd and Clark 1983). Nevertheless, we have demonstrated the delivery of a membrane-permeable fluorescent dye to *H. schachtii* and *M. hapla* male gonad, and the delivery of oligonucleotides to *M. hapla* male gonads. It should be emphasised that while this is possible, it is not routine, and most cells in the male *M. hapla* gonad do not spontaneously uptake even these short oligonucleotides. Recent demonstrations in animal parasitic nematodes have shown that the inclusion of lipofectamine in the injection mix, and subsequent injection into the gonad of *S. stercoralis* enabled CRISPR-Cas9 genome editing throughout the body (Adams et al. 2019). Our results suggest that including lipofectamine in the injection mix with fluorescently tagged oligonucleotides may increase the intercellular spread of the fluorescence, but we find no clear increase in cellular uptake in these species and at the concentrations used. It is possible that most cells in the gonad are too far differentiated to be receptive to cargo.

Despite their immature germline, the only other life stage of the sedentary endoparasites that is accessible to manipulation would be the J2. We explored whether lipofectamine could be used to deliver cargo to J2s by encapsulating mRNA encoding reporter genes in liposomes and using octopamine to stimulate nematode ingestion. Using this approach, we were able to demonstrate, for the first time, expression of exogenous mRNAs encoding fluorescent or bioluminescent proteins in a plant-parasitic nematode. Both fluorescence and bioluminescence are extremely sensitive methods of detection. While clearly above the background autofluorescence/noise thresholds for each technique, there is much that can be done to improve the signal to noise ratio, and thereby the utility of the approach. We expect major improvements in efficiency by optimising the reagents, for example the codon usage of mRNAs for *H. schachtii*, the inclusion of *H. schachtii* UTRs, and the use of lipofectamines specifically designed to encapsulate mRNAs. It is possible that modification of the mRNA may also improve stability and translation *in vivo* (Boo and Kim 2020). We also expect there is scope for improvement by optimising the experimental setup (e.g. time-of-soaking and time-of-detection, mRNA concentration, lipofectamine concentration etc.). Finally, we expect that addition of a nuclear localisation signal (in the case of fluorescent reporters) or epitope tag/s (in the case of bioluminescent reporters) may help to concentrate the signal, *in vivo* or *in vitro* respectively.

The ability to transiently express exogenous mRNAs in plant-parasitic nematodes is important and so worth optimising. Not only is it technically trivial and readily adopted without any specialised equipment, but it would also enable several experimental approaches that, until now, have been either impossible or prohibitively difficult for plant-parasitic nematology (e.g. *in vivo* protein-DNA interaction studies (ChIP seq) and *in vivo* protein-protein interaction studies (Co-IP, BiFC, FRET, etc.)). Most importantly, the observed increase in fluorescence in treated nematodes is throughout the body, not just in the digestive system of the nematode. This means that with sufficient optimisation of the technique, it may be possible to achieve expression of exogenous mRNA in the germ cell primordia of juvenile worms. Delivery in this way would avoid the difficulties of injecting cargo into male gonads, the subsequent uptake into male germ cells, the unknown challenges associated with mating females with males post injection, and in the case of the obligate parthenogens (e.g. *M. incognita*) the fact that males appear to not contribute genetically. Taken together, this transient expression system may also enable heritable genetic modification of plant-parasitic nematodes, through the delivery of mRNA encoding CRISPR-Cas variants to the germ cell primordia of juveniles.

We attempted to use a similar lipofection-based technique to deliver CRISPR-Cas9 protein to somatic cells of *H. schachtii* juveniles, but this was not successful. We initially tried to introduce a known edit by HDR, but the frequency of PCR-derived template switching was prohibitively high. We proceeded with an experiment designed to introduce a range of unknown edits by NHEJ in somatic cells of *H. schachtii* juveniles but were unable to consistently identify edited events. There are several potential explanations for these difficulties. The edited events are likely rare: if they occur at all, they may only be in a few cells per juvenile (PCR amplification of the locus does not selectively amplify edited events). Edits in individual cells are independent, and therefore may be different (even improving editing efficiency from 1 cell to 10 cells per nematode does not necessarily increase the signal, as each edit could be different). To the best of our knowledge, there are no known genetic modifications that would result in selectable dominant phenotypes in cyst or root knot nematode juveniles. Since the optimal temperature for *Streptococcus pyogenes* Cas9 (SpCas9) is around 37 °C, we envision that a heat-shock treatment may increase its efficiency (Xiang et al. 2017; LeBlanc et al. 2018). An additional possibility is to increase the accessibility of the enzyme to the target sequence by inducing an “open” chromatin state (Park et al. 2020; Chen et al. 2017). Finally, but perhaps most importantly, a successful CRISPR experiment requires several steps: 1) delivery of components into cells, 2) targeting a gene amenable to CRISPR, including guides that work *in vivo*, and 3) detection of possibly rare events - and failure at any one stage will result in failure. None of these steps have been established for any gene in plant-parasitic nematodes. It seems prudent to isolate and optimize as many of these challenges as possible. We anticipate that an optimisation of the protocol for lipofection-based delivery to juveniles described here will provide a route to rapidly overcome some of these challenges.

## Conclusions

Genetic modification of sedentary endoparasitic nematodes is an ongoing challenge. Delivery of cargo to their gonad by microinjection is difficult, but not impossible. Expressing exogenous mRNA throughout the juvenile body of *H. schachtii* is technically trivial, and potentially useful either on its own, or as a route to expedite the development of genetic modification protocol/s for sedentary endoparasitic nematodes.

## Materials and methods

### Biological material

Sand containing cysts of *Heterodera schachtii* was obtained from the Institute of Sugar Beet Research (IRS) in the Netherlands. Cysts were extracted from sand (by washing over 500 and 250-micron sieves with water) and collected in 50 mL falcon tubes. Juveniles were hatched by addition of 3 mM zinc-chloride and incubation at 20°C in the dark. Juveniles were collected at 2-3 day intervals, and stored in 0.01 % v/v Tween 20 in water at 4°C for up to 3 weeks until use.

To collect virgin males, sterile cysts were obtained from the University of Bonn (Germany), and maintained in sterile tissue culture on *Sinapis alba* (cv albatross) roots growing on KNOPs media at 20°C in the dark. Infected *Sinapis alba* roots were observed under a binocular microscope, and segments of root with differentiated J4s pre-emergence were collected, and placed in a 96-well plate. Each day, adult males that had emerged were removed and measured (for 0-1 dpe), or stored in the absence of females for n days until measured (for n days post emergence).

*Solanum lycopersicum* (cv Ailsa craig) roots infected with *M. hapla* were obtained from the University of Leeds (UK) and incubated in water in a petri dish to release males. Male nematodes were individually picked and stored in water at 4°C in the dark for up to 2 weeks until use.

### Microinjection of males

Nematodes were kept in ddH2O and washed with M9 buffer (www.wormbook.com) prior to injection. Animals were immobilised on 2-5 % w/v agarose pads which were prepared by placing a drop of hot agarose on a cover slip (22×50 mm) and rapidly placing another cover slip diagonally on top. Once the agarose had solidified, the coverslips on top were removed and agarose pads were dried at room temperature overnight. For immobilisation, a drop of Halocarbon oil 700 (Sigma H8898) was placed over the dried agar and washed worms were picked and placed in the oil using an eyelash pick. Microinjection was done using an Eppendorf Injectman and Femtojet set up on a Olympus inverted microscope with DIC prism. Worms were injected with either the Eppendorf pre-pulled Femtotips II or self pulled needles prepared from glass capillaries (Harvard Apparatus GC120F-15) using Sutter P-2000 instrument (Settings were Heat = 290, FIL = 4, VEL = 55, DEL = 225, PUL = 110). Injections were done using the highest pressure setting on the Femtojet (clean the needle function). Hoechst stain was used at 20 mM. Cy5.5 and FITC labelled oligos were injected at 100 μM. For lipofectamine injections 8 μl of 100 μM FITC oligo, 1.3 μl of CRISPRMax reagent and 0.7 μl H2O were mixed. *M. hapla* cuticle is very rigid and the needle angle had to be adjusted carefully to a near perpendicular angle to the animal body axis in order to prevent the needle tip from breaking.

### Microscopy

DIC images were taken using a Leica DMI6000 B inverted microscope equipped with a DIC prism. Hoechst, Cy5.5 and FITC images were taken using either a Leica SP5 or SP6 confocal imaging system. Live animals were placed on 2 % w/v fresh agarose pads with a 5 μl drop of 10 mM Levamisole and covered by a coverslip.

### Delivery of mRNA to plant-parasitic nematodes by lipofection

Capped and polyadenylated mRNAs encoding eGFP or Firefly luciferase were obtained from Ozbiosciences (Codon table not disclosed). To aid transfection of reporter mRNAs, two lipofection agents were used according to the manufacturer’s instructions. Approximately 15,000 - 20,000 J2 *H. schachtii* were soaked for 24 hours in 12 μg of mRNA, 12 % lipofectamine RNAIMAX or CRISPRMAX (Invitrogen), 100 mM octopamine (Thermo-Fisher), in a total volume of 30 μL (adjusted with nuclease-free M9 buffer). The negative control substituted the mRNA with an equal volume of M9 buffer.

### Detection of eGFP

Live or fixed (4% formaldehyde solution in PBS) transfected nematodes were transferred to a 76 x 26 mm microscope slide (Thermo scientific). The expression of eGFP was measured using a Leica SP5 confocal system mounted on a DM6 microscope equipped with an argon laser and photomultiplier tube (PMT) detectors. Z-stack images of the nematodes were collected with a 5 μm interval (ex 476 nm, em 508-513 nm, gain 714). Fluorescence difference between treated and control nematodes was visualized qualitatively. To provide a quantitative measure of fluorescence, the most in-focus optical section from the Z-stack was selected for each nematode manually. The image of each nematode was cropped out, inverted, their brightness, contrast, and intensity adjusted for both treatment and control (Brightness −17%, Contrast + 71%, Intensity - 27%), and the number of pixels per nematode that exceeded an empirically derived number of shades of variance (an integer between 0 and 255) from the background grey (Hexadecimal color code 0xD1D1D1) were counted using the PixelSearch function in a custom AutoIT script (https://github.com/sebastianevda/ColourCounter). The numerical value assigned to each nematode’s fluorescence allowed us to test the significance of the difference between the treatment and the control using an independent 2-group Mann-Whitney U test.

### Detection of luciferase

Luciferase expression was detected using the CLARIOstar plate reader (gain 4095). The supernatant was removed from the soaked nematodes. Each set of animals was resuspended in 240 μL nuclease-free M9 buffer, and distributed over 16 wells; eight positive (mRNA soaked) and eight negative controls (no mRNA). Using a CAPP^®^ 8-Channel Pipette (Starlab) 10 μl of 100 mM VivoGlo Luciferin (Promega) was added to the negatives and then to the mRNA soaked worms. The CLARIOstar plate reader was set up to vortex after each measurement at 300 rpm, and the plate was sealed with Corning^®^ microplate sealing tape (Sigma) and loaded into the machine and measured every 176 seconds. To quantitatively compare the dependency of luminescence as a function of time, the following model was fitted to the 16 time series (8 series for mRNA soaked, 8 for controls). The formula: Intensity = a + b * 2^(c*time). The model was fitted by the following R command: nls(y~a + b * 2^(c*time), start=list(a=1000, b = 1000, c = −0.0001)). The obtained half-lives (- 1 / c) in mRNA soaked is compared with the control half-lives using the independent 2-group Mann–Whitney U test.

### Delivery of CRISPR reagents to plant-parasitic nematodes by lipofection and analysis of target loci

Two-component guide RNAs (designed using CRISPOR (Haeussler et al. 2016) (Table S1)) were annealed by combining 3 μL Alt-R^®^ CRISPR-Cas9 tracrRNA, ATTO™ 550 with 4 μL Nuclease-free Duplex Buffer and finally 1.5 μL of Alt-R^®^ CRISPR-Cas9 crRNAs (Integrated DNA Technologies) and incubated at 95°C for 5 minutes and allowed to cool slowly to room temperature. The ribonucleoprotein complex was assembled *in vitro* by combining 3 μM gRNA either single guides (Synthego) with 25 μg of *Streptococcus pyogenes* 2xNLS-Cas9 protein (Synthego) or annealed crRNA:tracrRNA pair (IDT) with 25 μg of Alt-R^®^ *S. pyogenes* V3 Cas9 protein (IDT). After 5 minutes, the CRISPR-Cas9 ribonucleoprotein complexes were encapsulated in liposomes for 20 minutes at room temperature (3% v/v RNAIMAX (Invitrogen) or CRISPRMAX (Invitrogen) lipofectamine) and delivered to juveniles following essentially the same protocol as for mRNAs. In this case, octopamine (Sigma-Aldrich) was added to a final concentration of 50 mM, and the mixture was combined with 2,000 *H. schachtii* J2s and incubated at room temperature for 8 hours. The mixture was removed and the nematodes were washed 3 times in 200 μl of 0.01% v/v Tween 20 in sterile water. The DNA was extracted from transfected nematodes using the ChargeSwitch™ gDNA Mini Tissue Kit (Invitrogen) following the manufacturer’s instructions. Fragments containing the target site were amplified by PCR.

To assess template switching, a single stranded donor fragment (PAGE purified DNA oligo (IDT)) encoding a desired edit to the FAR-1-like gene of *H. schachtii* was co-encapsulated in liposomes with the ribonucleoprotein complex and relevant guide RNAs prior to delivery. Following DNA extraction, remaining ssDNA oligonucleotides were digested by the addition of 3 μL exonuclease 1 (NEB) to 20 μL extracted DNA, 3 μL exonuclease buffer (NEB) and 4 μL nuclease-free water (Ambion) and incubated at 37 °C for 15 minutes in an Eppendorf ThermoMixer^®^ (Eppendorf) followed by an incubation of 80 °C for 15 minutes to inactivate the exonuclease. The remaining DNA was used to amplify fragments of the target site by PCR and amplicons were purified using the Monarch^®^ PCR & DNA Cleanup Kit (NEB) following the manufacturer’s instructions and either sent for 250 bp paired end Illumina amplicon sequencing (Genewiz), or used as template in a second round of PCR with edit-specific primers and analysed by agarose gel electrophoresis. All template switching experiments were soaked for eight hours in the following three conditions: i) containing all components of the CRISPR-Cas reaction (termed positive), ii) all components of the CRISPR-Cas reaction with the exclusion of the relevant gRNAs (termed no-guide control), and iii) *H. schachtii* J2s without additional reagents (termed negative control).

To assess non-homologous end joining, ten *H. schachtii* genes were selected based on their expression at J2 (Pers. comm. Eves-van den Akker) and/or putative function assigned by sequence homology to *C. elegans*. Sequences of genes of interest are available in DRYAD repository XYZ. A total of 28 gRNAs over the ten genes were designed using CRISPOR (Haeussler et al. 2016) (Table S1). Transfection experiments were performed, largely as above, for 24 hours with the following two conditions: i) containing all components of the CRISPR-Cas reaction (termed positive), and ii) all components of the CRISPR-Cas reaction with the exclusion of the relevant gRNAs (termed negative). DNA was extracted from transfected nematodes and the region of interest amplified proof reading PCR (primers in Table S1). Amplicons were purified using the Monarch^®^ PCR & DNA Cleanup Kit (NEB) following the manufacturer’s instructions and sent for 250 bp paired end Illumina amplicon sequencing (Genewiz). Reads were trimmed (Phred-64) and overlapping pairs were re-capitulated into the amplified fragment using scripts designed for similar metagenetic analyses (Eves-van den Akker et al. 2015) (https://github.com/sebastianevda/SEvdA_metagen). Recapitulated fragments were further analysed for edits within the guide region using a set if custom shell and python scripts (https://github.com/OlafKranse/Selective-analyses-of-areas-of-interest-for-next-generation-sequencing). In brief: the most common amplicon was set as reference; the regions targeted by the guide RNAs in this reference were located; unique reads are aligned individually to the new reference; the sequences within the guide location (6 bp upstream of PAM) are compared; if there is a difference in sequence, a record is made containing which type of difference (e.g. SNP or INDEL) and the number of occurrences of that specific mutation.

## Data availability

NGS reads deposited under ENA accession PRJEB39266. Scripts available at github repositories https://github.com/sebastianevda/ColourCounter and https://github.com/OlafKranse/Selective-analyses-of-areas-of-interest-for-next-generation-sequencing. Gene sequences available at DRYAD under accession doi:10.5061/dryad.r4xgxd296.

## Acknowledgements

Work on plant-parasitic nematodes at the University of Cambridge is supported by DEFRA licence 125034/359149/3, and was funded by BBSRC grants BB/R011311/1, BB/N021908/1, and BB/S006397/1, a Synthego Genome Engineer grant, and a Genewiz grant. AA was supported by a Wellcome/Newton trust Institutional Strategic Support Fund grant and a UKRI Future Leaders Fellowship MR/S033769/1. CJL was supported by BBSRC grant BB/N016866/1. JJ receives funding from the Scottish Government Rural and Environmental Science and Analytical Services division. TRM and TJB were supported by grants from the Iowa Soybean Association and by State of Iowa and Hatch funds.

---

**Table S1.**
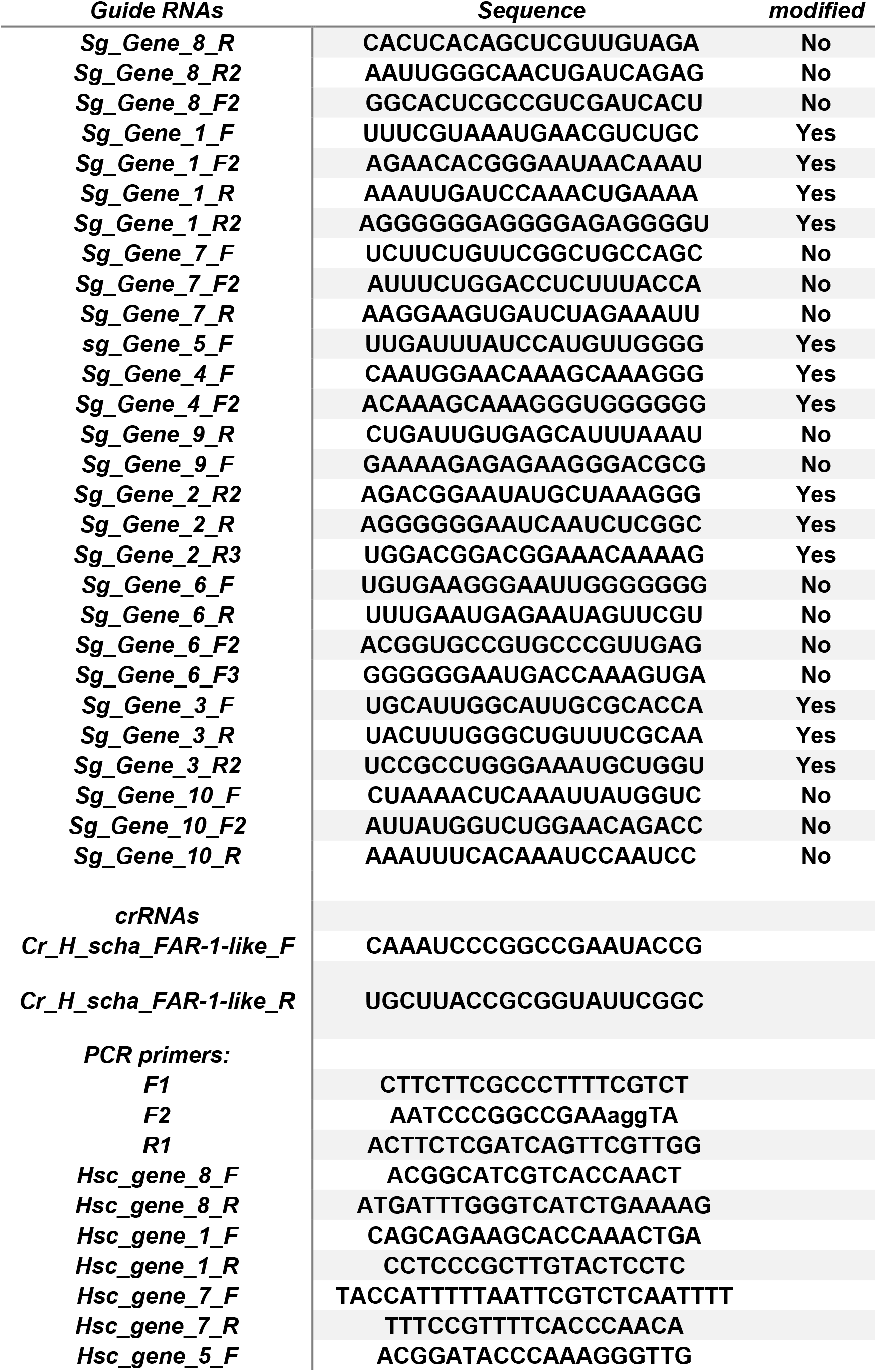
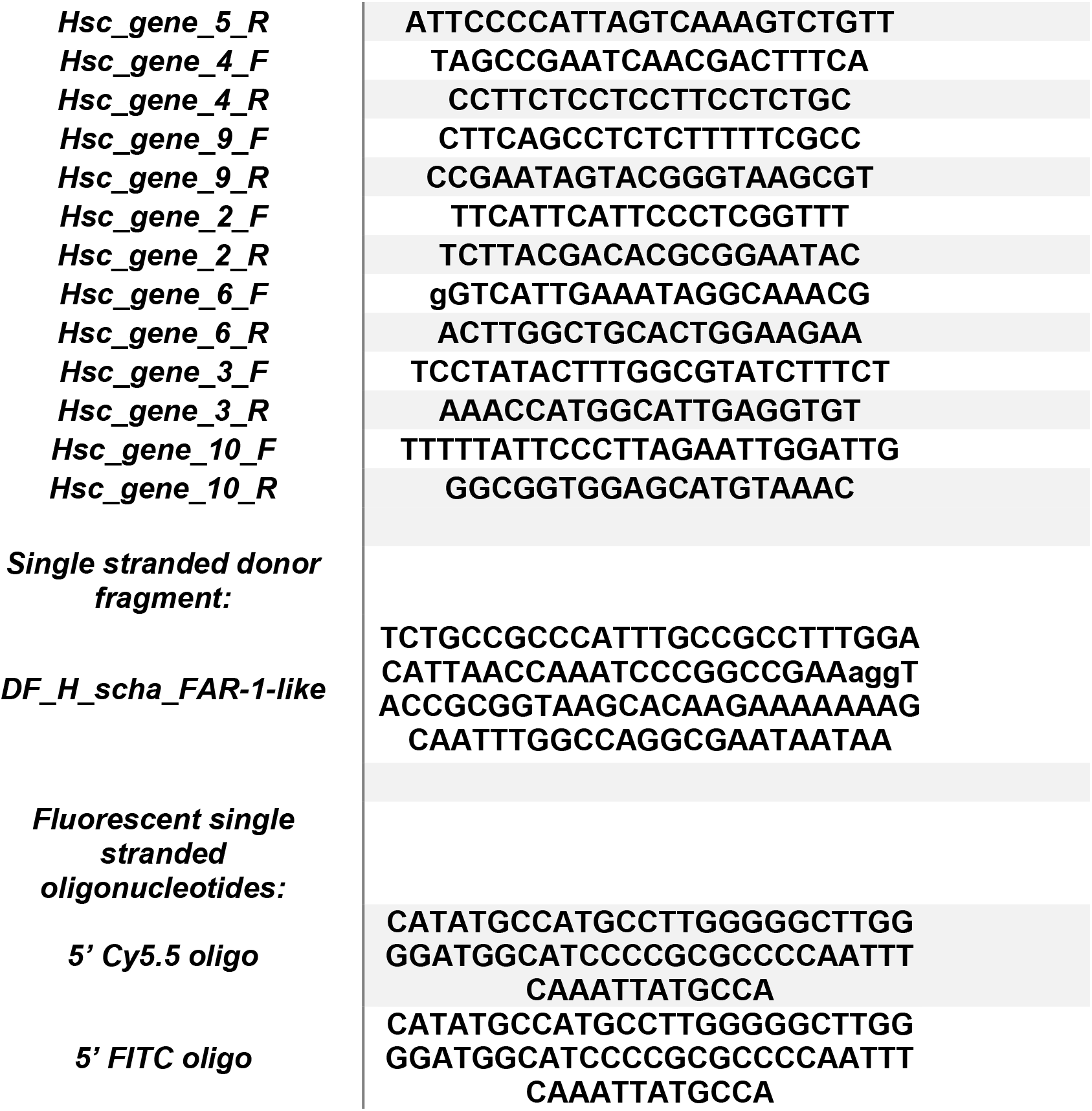
Oligonucleotides.

## Notes

https://doi.org/10.5061/dryad.r4xgxd296

https://github.com/sebastianevda/ColourCounter

https://github.com/OlafKranse/Selective-analyses-of-areas-of-interest-for-next-generation-sequencing

https://www.ebi.ac.uk/ena/data/view/PRJEB39266

### Competing Interest Statement

The authors have declared no competing interest.

